# Programming Metabolic Dependency in Synthetic Cells Under Resource Scarcity

**DOI:** 10.64898/2026.07.01.735941

**Authors:** Basak Velioglu Ulubas, Orion Venero, David Garenne, Vincent Noireaux, Aaron E. Engelhart, Katarzyna P. Adamala

## Abstract

Engineering synthetic cells where metabolism directly controls gene expression is one of the greatest challenges in synthetic biology. This coupling between metabolic activity and protein synthesis is also thought to have been a vital step in the evolution of the earliest cellular life. This fundamental process is essential for developing minimal, self-sustaining cells, both engineered for biotechnology applications and as models explaining the origins of life. Here, we present a programmable cell-free platform that links metabolic activity to translation under defined resource limitations. Using an engineered amino-acid-dependent cell-free translation system, we introduced a tunable metabolic bottleneck (depleting tyrosine). This enabled imposing a controlled metabolic constraint on protein synthesis. To alleviate this constraint, we then incorporated phenylalanine hydroxylase (PAH) as a minimal module for tyrosine synthesis. The PAH-driven tyrosine synthesis established a system in which protein expression is directly controlled by amino acid biosynthesis. This relationship was recapitulated in liposome compartments, resulting in three distinct synthetic cell populations. The metabolically active population had significantly higher fitness in protein production. Overall, this work establishes an experimentally tractable platform to investigate how primitive cells may have evolved internal metabolic capabilities, and it represents a foundational step toward constructing more autonomous and self-regulating synthetic cells.

## Introduction

Synthetic cells provide a powerful platform for reconstructing minimal biological systems, bypassing the complexity of natural cellular environments. Various biological functions can be incorporated into synthetic cells to make them more lifelike and increasingly useful in biotechnological applications, such as biomedical diagnostics and biosensors ^1,2,3^. Consensus on life-like properties suggests that protocells must preserve and utilize genetic information through the central dogma, thereby requiring a processable informational genome ^4,5^. Second, protocells must possess metabolic capabilities to generate energy and synthesize essential molecular building blocks^4,6^. Third, these processes must operate within an organized structure that enables their coordinated functioning, which can be achieved through compartmentalization ^4,7^. Although several studies have successfully demonstrated that core biological modules can coexist in synthetic cells, they rarely establish direct functional coupling between metabolism and gene expression^8^. In natural cells, however, metabolic flux strongly constrains protein synthesis by determining the availability of essential metabolites. Recreating this dependency represents a critical step toward engineering autonomous synthetic cells.

Insights from early cellular evolution may guide for achieving this integration and its importance in self-sustainability. Early protocells are widely hypothesized to have depended heavily on environmental resources that were limited in availability and diversity. Therefore, within this ancestral community of primitive cells, lineages that evolved metabolic pathways capable of synthesizing essential molecules internally - rather than depending solely on environmental supply - would have gained a selective advantage.

The emergence of internal metabolic pathways may therefore have been a key step toward increasingly autonomous, more robust cellular systems. In this study, we leverage this concept to engineer synthetic cells in which metabolic activity directly influences gene expression.

To establish this link, we construct a model system that couples a minimal metabolic module to two different cell-free expression systems, TxTl and PURE. TxTl is a widely adopted, robust expression system that utilizes whole-cell extracts ^9,10^. Protein Synthesis Using Recombinant Elements (PURE), on the other hand, is a translation system composed of purified bacterial ribosomes and other proteins, offering a well-defined mixture with known components and concentrations ^11^. The metabolic module is run by phenylalanine-4-hydroxylase (PAH), an enzyme key to tyrosine metabolism that converts phenylalanine into tyrosine with tetrahydropterin (BH□), non-heme Fe² cofactors, and oxygen^12,13^. We selected *Chromobacterium violaceum* PAH (cPAH) due to its extensive study and the absence of post-translational modifications typical in mammalian enzymes ^12,14^. For synthetic cell experiments, we used phospholipid liposomes to encapsulate translation and metabolic reactions.

Overall, we demonstrate a genetically encoded coupling between metabolism and the central dogma within synthetic cells. This approach has the potential to foster the development of more robust, controllable cell-free systems and facilitate the creation of complex, dynamic artificial biological systems capable of autonomous behaviors.

## Results and Discussion

### Amino acid scarcity was simulated in cell-free expression via dialysis

To develop an amino acid resource scarcity expression system, we identified key amino acids vital for cell-free translation (Fig. 1a). For our experiments, we selected GFP reporter^15^, whose chromophore is a tripeptide composed of serine, tyrosine, and glycine (Ser65-Tyr66-Gly67)^16^. We hypothesized that limiting the availability of any of these amino acids would effectively prevent GFP fluorescence. Because lysate expression systems contain all components of *E. coli* cytoplasm^17^, the cellular extract likely contains residual amino acids. This allows some level of expression without external amino acid supplementation. To remove those endogenous amino acids, we dialyzed *E. coli* extract before cell-free expression (Fig. 1a), making the system reliant on externally supplied amino acids (and all other small molecule substrates).

**Fig. 1.**
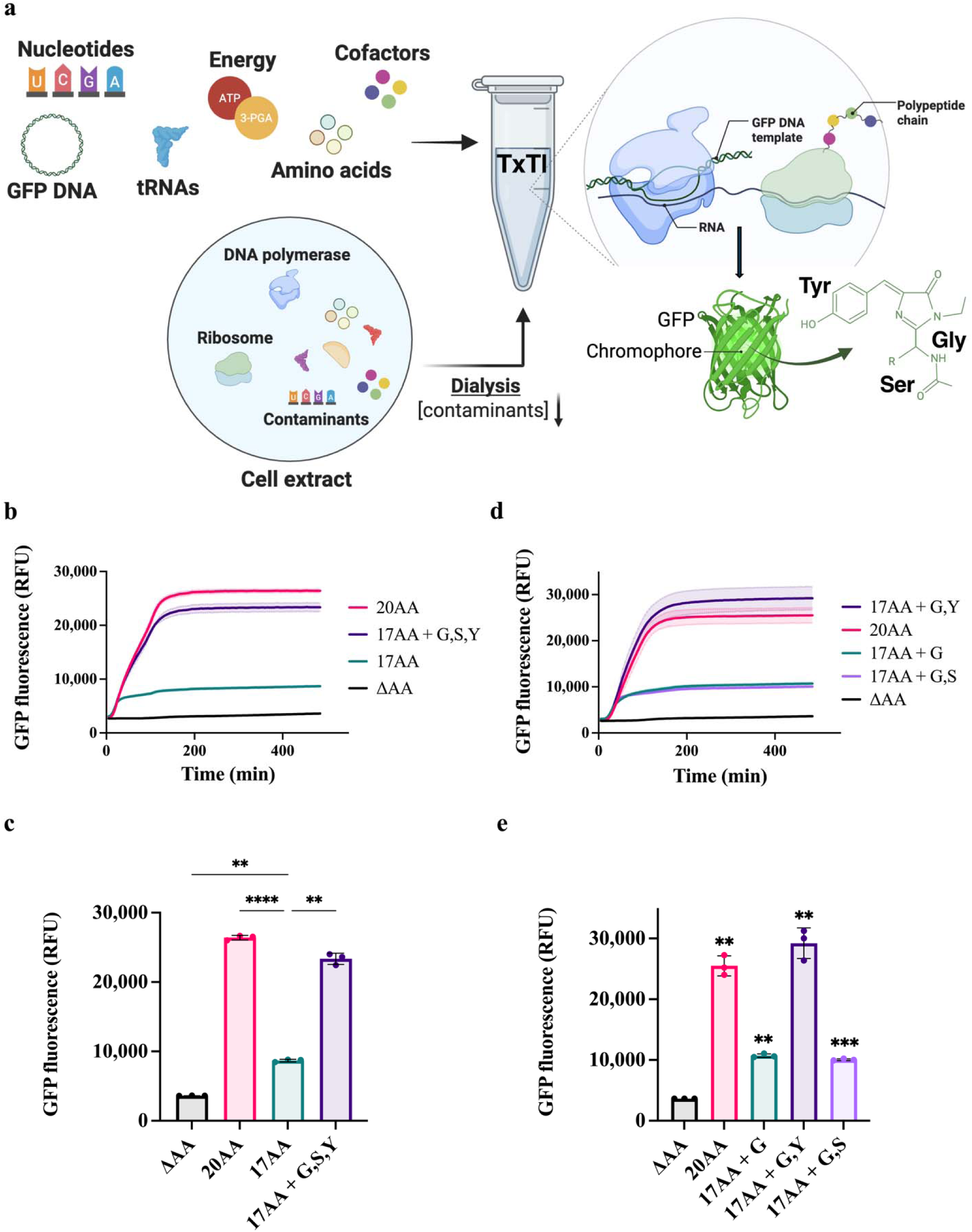
Resource scarcity for TxTl was simulated through amino acid depletion. **a.** General scheme of the removal of amino acids carried over with the cellular extract, and validating GFP expression under amino acid depletion conditions. Cell extract was dialyzed to eliminate amino acids. TxTl was performed using dialyzed cell extract, supplemented by either a limited amino acid set or a complete set of 20 amino acids. **b.** Real-time fluorescence assay showing the GFP expression without external Gly, Ser, and Tyr. **c.** End point analysis showed that removing these amino acids altogether decreases GFP expression significantly. **d.** Real-time fluorescence assay showing the GFP expression using dialyzed TxTl without external Gly, Ser, and Tyr separately. **e.** The end point analysis showed that the internal Gly and Ser concentrations in the cell extract were sufficient even after dialysis, but extracellular Tyr was required for efficient expression. Statistical significance was determined using ordinary one-way ANOVA. Error bars indicate standard deviation (SD). Data is expressed as the mean + SD.

Using the dialyzed system, we initially examined how removing Gly, Ser, and Tyr from the external amino acid supply affected GFP expression. All experiments were performed using 15 nM GFP plasmid, and the GFP fluorescent signal was detected using a Bio-Rad qPCR system under SYBR filter. We found that eliminating these three amino acids significantly decreased GFP expression, and expression could be restored by reintroducing them via the external supply (Fig. 1b and 1c).

Next, we focused on the individual effects of these amino acids on GFP expression, and discovered that Tyr, but not Ser or Gly, significantly lowered GFP translation efficiency (Supplementary Fig. 1a and 1b, and Fig.1d and 1e). We hypothesize that this is because Gly and Ser are abundantly present in the cell extract, regardless of dialysis, while the concentration of Tyr can be significantly reduced through dialysis, making its availability mostly dependent on external supply. And since Gly and Ser are highly abundant, their levels might be maintained by protein degradation within the TxTl system, thereby reducing their external requirements^18^. Consistent with this, even in the absence of external Tyr supply, residual background expression was observed, likely due to Tyr recycling via protein degradation (Fig. 1d and 1e).

Next, we investigated the concentration of externally supplied Tyr to rescue GFP translation. We tested Tyr concentrations, ranging from 2 mM to 0.03 mM, and we demonstrated that GFP expression could be restored with Tyr concentrations as low as 0.03 mM, compared to the standard amino acid supply of 2 mM (Supplementary Fig. 1c and 1d). Overall, these results demonstrated that selective pressure can be created through the dialysis of TxTl to deprive the system of Tyr amino acid, and external Tyr supply can be used to regulate the cell-free translation activity of GFP.

### A metabolism-dependent expression system was developed utilizing phenylalanine-4-hydroxylase

We chose the PAH enzyme from *C. violaceum* for the biosynthesis of the amino acid Tyr. This enzyme catalyzes the oxidation of Phe to Tyr under oxygen-rich conditions, utilizing the cofactor 6,7-dimethyl-5,6,7,8-tetrahydropterin (DMPH□) and Fe^2+^ ^12^(Fig. 2a). First, we expressed PAH in *E.coli*, purified, and characterized its activity (Supplementary Fig. 2). The enzymatic reaction was optimized in vitro, using UV-vis spectroscopy to detect Tyr product. We found that in vitro optimized PAH reaction (2 mM Phe, 0.5 mM DMPH□, 1.0 µM PAH, 4 mM DTT, 6 µM FeSO□ in 0.1 M HEPES (pH 7.6) yielded a Tyr concentration of 0.13 mM, surpassing the threshold (0.03mM) needed for efficient GFP expression with Δtyr-TxTl (Supplementary Fig. 1c and 1d). We calculated Michaelis-Menten parameters based on Tyr standards (Supplementary Fig. 2b, 2c, and 2d). These parameters allowed us to estimate the time required to synthesize 0.03 mM Tyr. With a 2 mM Phe supply, we calculated that PAH takes 5 minutes to synthesize 0.03 mM Tyr at 37°C. However, this should be considered an estimate, as a simultaneous TxTl reaction might interfere with the PAH enzymatic components and affect kinetic parameters. Still, we hypothesized that the PAH reaction is sufficient to generate Tyr required for GFP translation, depending on these results.

**Fig. 2.**
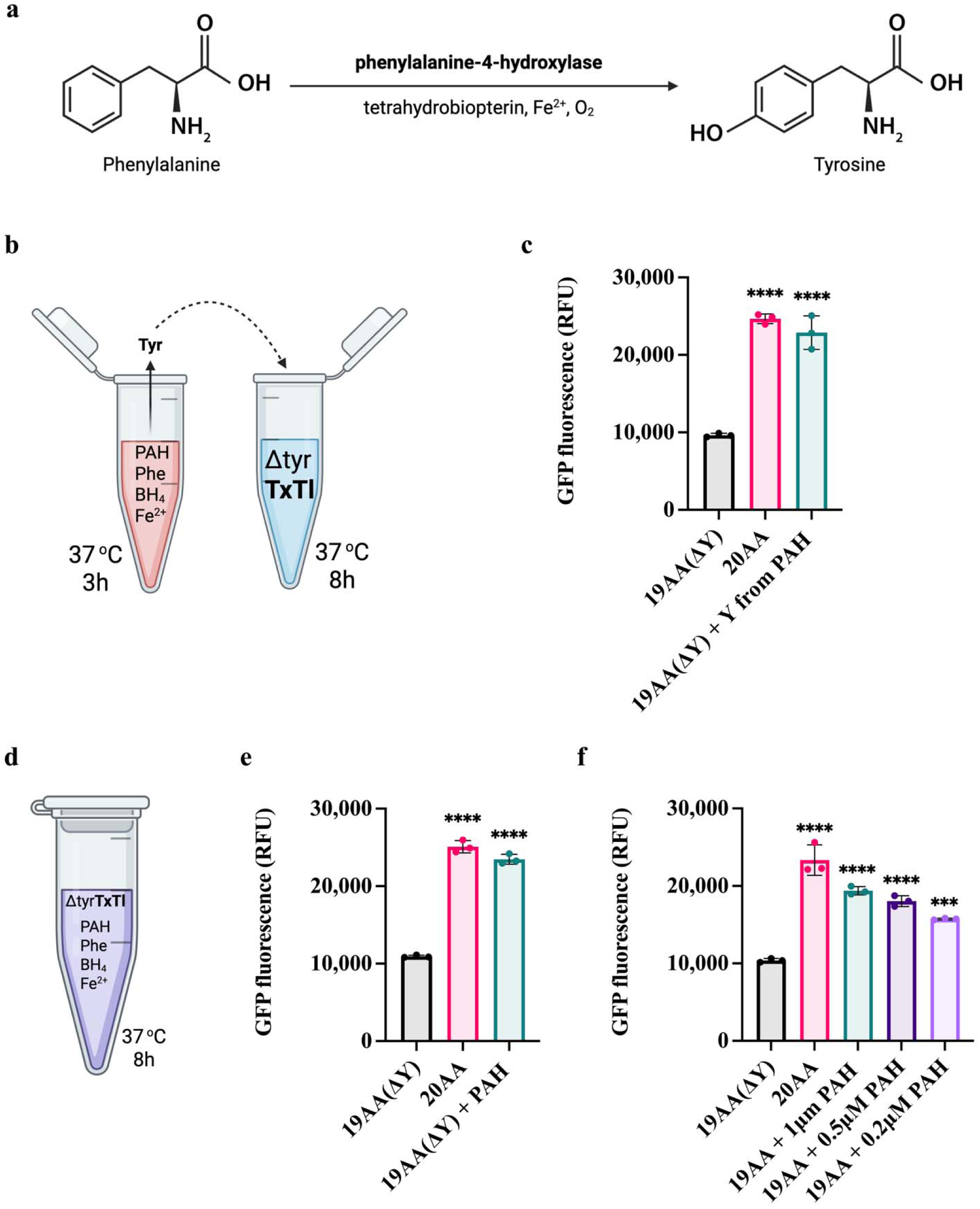
The TxTl and PAH reactions were coupled in vitro. **a.** Schematic representation of the enzymatic synthesis of Tyr using the PAH enzyme. Tyr synthesis requires the substrates Phe and BH_4_, and the cofactor Fe^2+^ under oxygen-rich conditions. **b.** Schematic representation of the experimental setup in which components required for Tyr synthesis were preincubated for 3 hours and then served as the Tyr source for Δtyr-TxTl. **c.** End-point analysis showed that the Tyr produced via the PAH reaction effectively rescued GFP expression in Δtyr-TxTl. **d.** Schematic representation of the experimental setup in which the PAH reaction synthesizing Tyr and the Δtyr-TxTl expressing GFP were initiated simultaneously. **e.** End point analysis showed the compatibility of the metabolic reaction with cell-free expression. PAH reaction can operate simultaneously with TxTl expression. **f.** The coupling of GFP expression with the PAH reaction was tested using different PAH enzyme concentrations. End point analysis showed that decreasing PAH enzyme concentration significantly reduced GFP expression in Δtyr-TxTl. Statistical significance was determined using ordinary one-way ANOVA. Error bars indicate standard deviation (SD). Data is expressed as the mean ±SD.

After confirming that the PAH reaction is a feasible metabolic pathway to rescuing Tyr-depleted translation, we assessed the effects of PAH reaction components on GFP expression. All experiments were performed using 15 nM GFP plasmid. After a 3-hour in vitro Tyr synthesis at 37°C, the PAH reaction mixture was used as a Tyr source for GFP expression (Fig. 2b). All TxTl samples included PAH reaction components, but only the test sample contained active PAH enzyme. Results showed that the preincubated PAH reaction mixture could rescue GFP translation, without inhibitory effects on GFP synthesis (Fig. 2c). In a subsequent experiment, we initiated simultaneous reactions in a single tube to evaluate compatibility between TxTl and PAH reactions (Fig. 2d). We demonstrated efficient coupling of GFP expression with PAH reaction (Fig. 2e). Finally, we systematically varied PAH concentrations to assess the regulatory capacity of PAH. We observed that GFP expression drops noticeably as enzyme concentration decreases (Fig. 2f). This demonstrates that adjusting only the PAH concentration can directly regulate protein expression.

### Genetically encoded amino acid metabolism coupled with cell-free expression

Having established PAH as a viable route to rescue Tyr deficiency in TxTl, we moved on to establish the metabolic rescue reaction on the genetic level (expressing PAH in the same reaction as it is rescuing). Initially, we evaluated PAH expression in Δtyr-TXTL. To optimize PAH synthesis, we made several modifications to the TxTl protocol. First, the reaction temperature was reduced to 30 °C, and the incubation time was extended to 16 hours to minimize PAH aggregation ^19^. Second, the backbone vector was optimized to increase TxTl efficiency (introducing a more optimal Shine-Dalgarno sequence and start codon spacing, and lower GC content that reduces mRNA hairpin formation). Thirdly, we switch to the sfGFP reporter to boost the overall fluorescence signal. As anticipated, all those changes resulted in a stronger fluorescence signal.

Our results in the previous sections indicate that a trace amount of Tyr is present in the Δtyr-TxTl, as evidenced by the background signal. This allows the PAH enzyme to be expressed with Δtyr-TxTl without external Tyr supply. Background-level expression with 19AA(Δtyr) shows approximately one-third the efficiency of 20AA-TxTl (Supplementary Fig. 1c and 1d). In the TxTl system we work with, TxTl 2.0, the maximum protein yield we could expect from 20AA-TxTl is approximately 2□mg/mL in batch reactions ^20^. Assuming similar yields, the background reaction could produce approximately 0.67 mg/mL of protein. Using Equation (1), we calculated PAH expressed using Δtyr-TxTl to be 20□μM - twenty times the amount needed for restoring full sfGFP expression (comparable to 1□μM purified PAH enzyme) (Fig. 2f). Based on these calculations, we anticipated that we could synthesize sufficient PAH using the trace amount of Tyr present in Δtyr-TxTl. To test this, we expressed 5□nM PAH plasmid and performed Western blot analysis. As expected, PAH synthesis with 19AA was as efficient as that with 20AA (Fig. 3a).

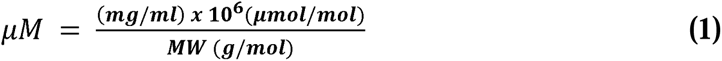

**Fig. 3.**
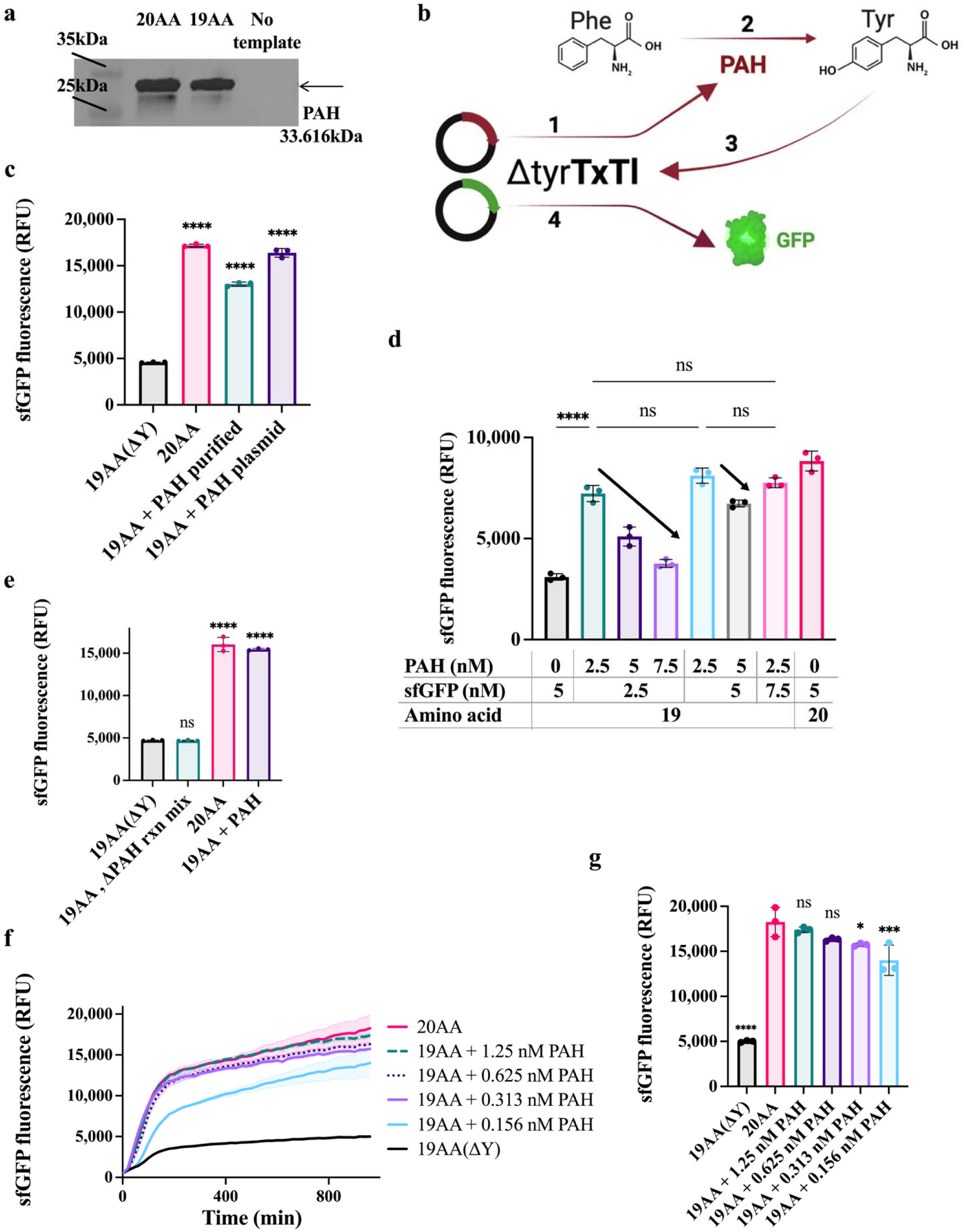
Amino acid metabolism was genetically incorporated into cell-free expression. **a.** Western blot analysis of PAH after expression via TxTl with 19AA and 20AA. **b.** Schematic illustration of PAH reaction-dependent TxTl expression. The PAH plasmid can be expressed using Tyr-deficient TXTL due to trace Tyr content in the extract; however, GFP expression remained dependent on the PAH reaction. **c.** The efficiency of sfGFP expression was tested using both 1 µM purified PAH and 5nM PAH plasmid. End point analysis showed that plasmid PAH rescued sfGFP expression as effectively as the purified enzyme did. **d.** The concentrations of PAH and sfGFP template were optimized to maximize sfGFP expression. End point analysis showed that increasing PAH concentration hindered sfGFP expression due to limited TXTL resources. **e.** Metabolism and expression coupling were enhanced further, excluding Phe substrate in the PAH activity assay. The Phe supply for the Δtyr-TxTl reaction was shared between PAH and sfGFP expression, as well as for the PAH-catalyzed reaction. End point analysis showed that PAH reaction mix does not have any effect on the background expression. Overall background level dropped with the exclusion of additional Phe from the PAH activity assay. **f.** Further decreasing PAH plasmid concentrations revealed the threshold for dropping sfGFP expression in real-time fluorescence reading. **g.** End point analysis shows that the fluorescence begins to decline at and below 0.313 nM PAH plasmid supplementation. Statistical significance was determined using ordinary one-way ANOVA. Error bars indicate standard deviation (SD). Data is expressed as the mean ± SD.

Next, we tested the coupled system, with PAH plasmid integrated in Δtyr-TxTl-based sfGFP expression (Fig. 3b). We performed the Δtyr-TxTl reaction with 5nM PAH and 5nM sfGFP. We showed that active PAH was expressed from the plasmid, using Δtyr-TxTl at 30 °C, and concurrently it rescued sfGFP expression at least as effectively as purified PAH (Fig. 3c).

We then examined this genetic coupling in more detail by varying the concentration of the PAH plasmid, measuring its effect on sfGFP expression levels. Surprisingly, we found that increasing PAH plasmid concentration did not enhance sfGFP expression; instead, it significantly reduced it. We hypothesize that this decrease is due to the competition for shared components in TxTl between sfGFP and PAH.

Overall, the highest sfGFP fluorescence was achieved at PAH concentrations equal to or below those of sfGFP, specifically at 5□nM sfGFP with 2.5□nM PAH, 2.5nM sfGFP with 2.5□nM PAH, and 7.5□nM sfGFP with 2.5□nM PAH (Fig. 3d). These results indicate that the PAH expression must be tuned to remain in equilibrium with expression of other proteins in the same reaction. Otherwise, PAH consumes critical resources needed for sfGFP expression. We hypothesize that employing a dialysis system to replenish energy sources could help overcome those resource limitations in TxTl. Since we had already achieved 20□AA-level activity using the PAH metabolic reaction in Δtyr-TxTl, we considered switching experimental systems unnecessary. In the future, this approach could also be extended by incorporating semi-continuous energy supplementation to further enhance overall expression efficiency ^21^.

To further strengthen metabolic coupling, we removed the 2□mM Phe required for PAH activity, thereby making PAH activity dependent on Phe supply in Δtyr-TxTl. We observed that PAH reactions can function efficiently without an additional Phe supply (Fig. 3e). This shared substrate strengthens the metabolic coupling between PAH and GFP (the biosynthetic enzyme and the reporter). Additionally, a slight decrease in background was observed at lower Phe concentration.

We anticipated that Phe, when used in excess in the Δtyr-TxTl system, could generate a background signal through occasional misincorporation at Tyr codons. Because Phe and Tyr differ by one para-hydroxyl group, TyrRS’s selectivity against noncognate Phe drops under tyrosine-depleted conditions, aligning with reports of TyrRS mischarging tRNA^Tyr with Phe at rates up to 0.7% under tyrosine limitation^26^. On the other hand, the PAH reaction mix itself did not increase the background signal.

Finally, we determined the minimum PAH plasmid concentration required to achieve full sfGFP fluorescence. We found that 0.625nM PAH is sufficient for complete recovery of sfGFP fluorescence, and at 0.313nM, a drop in efficiency begins to appear (Fig. 3f and 3g). Consistent with our earlier findings, these results indicate that sfGFP expression can be genetically controlled by adjusting the plasmid concentration of the enzyme in the biosynthetic metabolism.

### Universality of metabolism-dependent expression was demonstrated using PURE

Encouraged by our results, we aimed to demonstrate that metabolism-dependent expression systems persist across various CFPS platforms. We initially assessed the expression of the PAH enzyme using PURExpress with a complete set of 20 amino acids and excluding Tyr. Following expression, Western blot analysis revealed that PAH could be successfully produced with the full amino acid mixture; however, without external Tyr supplementation no PAH band was detected (Fig. 4a). Interestingly, even without external Tyr, PAH was detectable if the PAH activity mixture was supplied during its expression (Fig. 4a). This finding suggests that, even in very small amounts, PAH was expressed with Tyr-excluded PURE and capable of synthesizing Tyr. This Tyr, synthesized by PAH, in turn, appears to enhance PAH expression, making it detectable via Western blot.

**Fig. 4.**
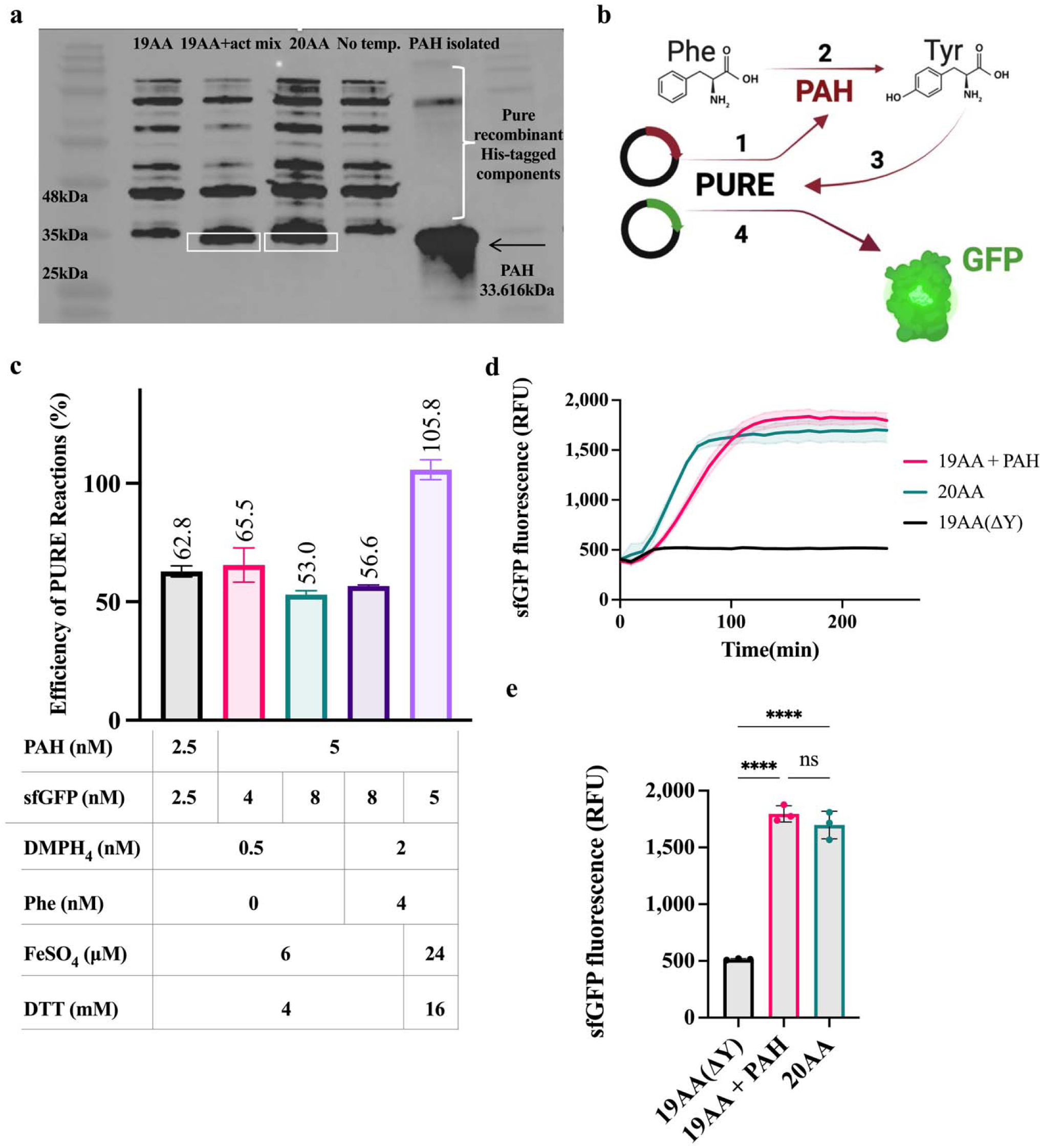
Amino acid metabolism was genetically incorporated into PURE. **a.** Western blot analysis of PAH after expression via PURE. PAH can be expressed using either 20AA or 19AA, along with PAH activity supplements through PURE. **b.** Schematic illustration of PAH reaction-dependent PURE expression. **c.** Efficiency of the PURE expression system with various plasmids and PAH activity supplement formulations. It summarizes the step-by-step optimization process, accounting for endpoint readings in efficiency calculations. Efficiencies of GFP expression using 19AA and PAH reaction were calculated based on the average GFP fluorescence using 20AA. The final bar showing more than 100% efficiency is due to the reader defect. **d.** Fully optimized metabolism-dependent PURE system analyzed with real-time fluorescence analysis. **e.** End point analysis showed that GFP expression in PURE was fully recovered through the PAH reaction. Statistical significance was determined using ordinary one-way ANOVA. Error bars indicate standard deviation (SD). Data is expressed as the mean ± SD.

Based on these results, we aimed to assess PAH activity through real-time fluorescence assay, hypothesizing that even minimal PAH expression from 19 amino acids could initiate tyrosine synthesis, allowing the enzyme to rescue sfGFP expression (Fig. 4b). Thus, we began testing a PAH - sfGFP coupled system, maintaining identical plasmid and PAH reaction component concentrations as in Δtyr-TxTl. We initially observed that optimized plasmid concentrations for Δtyr-TxTl could not be directly applied to the PURE system. Specifically, we achieved a 62.8% sfGFP expression efficiency with 2.5 nM sfGFP and 2.5 nM PAH (Supplementary Fig. 3a and Fig. 4c). We hypothesized that this discrepancy stems from the TxTl system’s robust inherent metabolic pathways for energy regeneration and the presence of contaminants such as amino acids and cofactors, which compensate for source requirements. Next, we tried increasing plasmid concentrations to boost overall expression, using 5 nM sfGFP and 4 nM PAH. Unfortunately, this did not enhance sfGFP expression as well (Supplementary Fig. 3b and Fig. 4c). Following that, we raised PAH concentration from 4 nM to 8 nM while keeping sfGFP at 5 nM; however, instead of an increase, we observed a drop in efficiency (Supplementary Fig. 3c and Fig. 4c). This result was likely caused by the excess PAH plasmid consuming the PURE sources, which is consistent with the expression using TxTl. This indicates that source competition in two plasmid systems was observed regardless of the purity of the expression systems.

Subsequently, we optimized the PAH activity assay rather than the PURE system itself. Keeping plasmid concentrations at 8 mM PAH and 5nM sfGFP, we increased the Phe substrate level to 4 mM and the DMPH_4_ level to 2 mM. This modification resulted in only a 56.6% sfGFP yield (Supplementary Fig. 3d and Fig. 4c). Our final optimization involved maintaining the same plasmid and substrate concentrations and increasing the concentrations of the cofactor, ferrous sulfate (24 μM), and DTT (16 mM). This approach finally enabled fully efficient sfGFP expression in the PURE system (Fig. 4c, Fig. 4d, and 4e). Additionally, the same result reveals a lag in sfGFP fluorescence (Fig. 4d), indicating that PURE sources are shared between PAH and sfGFP expression, and that sfGFP translation alone cannot proceed efficiently without PAH activity. This confirms that the Tyr-excluded PURE system relies entirely on PAH expression and PAH-dependent tyrosine synthesis for optimal sfGFP production. Notably, the PURE system showed no background expression with 19 external amino acid supply, suggesting the absence of Tyr contamination that could otherwise induce a base-level sfGFP signal.

Overall, these findings probe the boundaries of the ability to regulate the PURE expression system through metabolic feedback.

### Metabolic dependency was built into synthetic cells

After demonstrating that amino acid metabolism can be integrated into expression systems, we sought to extend this methodology to synthetic cells as a model for the emergence of essential metabolic reactions under source-scarce conditions. We hypothesized that, in liposomes, PAH enzymes could be expressed via Δtyr-TxTl and, using substrates and cofactors, synthesize Tyr, which is vital for sfGFP expression.

To test this hypothesis, we prepared liposomes using an emulsion transfer method as previously described^30^. Initial tests involved encapsulating a TxTl system containing only sfGFP with 20 amino acids to assess encapsulation efficiency and readout capacity. After 16 hours of TxTl reaction at 30°C, liposomes bearing the sfGFP template showed 4.2 times higher fluorescence than the no-template control (Supplementary Fig. 4a and 4b). On the other hand, without encapsulation, the same TxTl reaction yielded a 20-fold sfGFP signal compared to the no-template control (Supplementary Fig. 4c, 4d, and 4e), indicating a 4.8-fold reduction in signal detection due to light scattering within the liposomes. Consequently, we estimated that the difference in sfGFP signal between PAH Δtyr-TxTl and the background Δtyr-TxTl would be minimal in liposomes in a fluorescent reader population-based analysis. As expected, liposomes with PAH exhibited only a 1.2-fold increase in sfGFP fluorescence compared to the background control (Supplementary Fig. 4f and 4g).

To facilitate more detailed visual analysis, we conducted fluorescent microscopy on three different liposome populations: population 1 - no template (Δtyr-TXTL), population 2 - background (sfGFP Δtyr-TXTL), and population 3 - metabolically active (sfGFP and PAH Δtyr-TxTl) (Fig. 5a). We analyzed 30 liposomes per population, based on their radius and mean fluorescence intensity and found that metabolically active population has greater fluorescence signal compared to background and no template control (Fig. 5b). Additionally, the liposome distribution analysis showed that larger liposomes have a more pronounced fluorescence signal especially in population 3. We anticipated that the size effect could arise from greater encapsulation efficiency of cofactors and substrates required for PAH activity. The difference between fluorescence intensities was also validated through region of interest (ROI) analysis on raw microscopy images of the three replicates per population, by examining 10 positive liposomes and 3 background regions in each image. ROI analysis showed that population 3 has significantly greater GFP fluorescence compared to populations 2 and 1 (Fig. 5c).

**Fig. 5.**
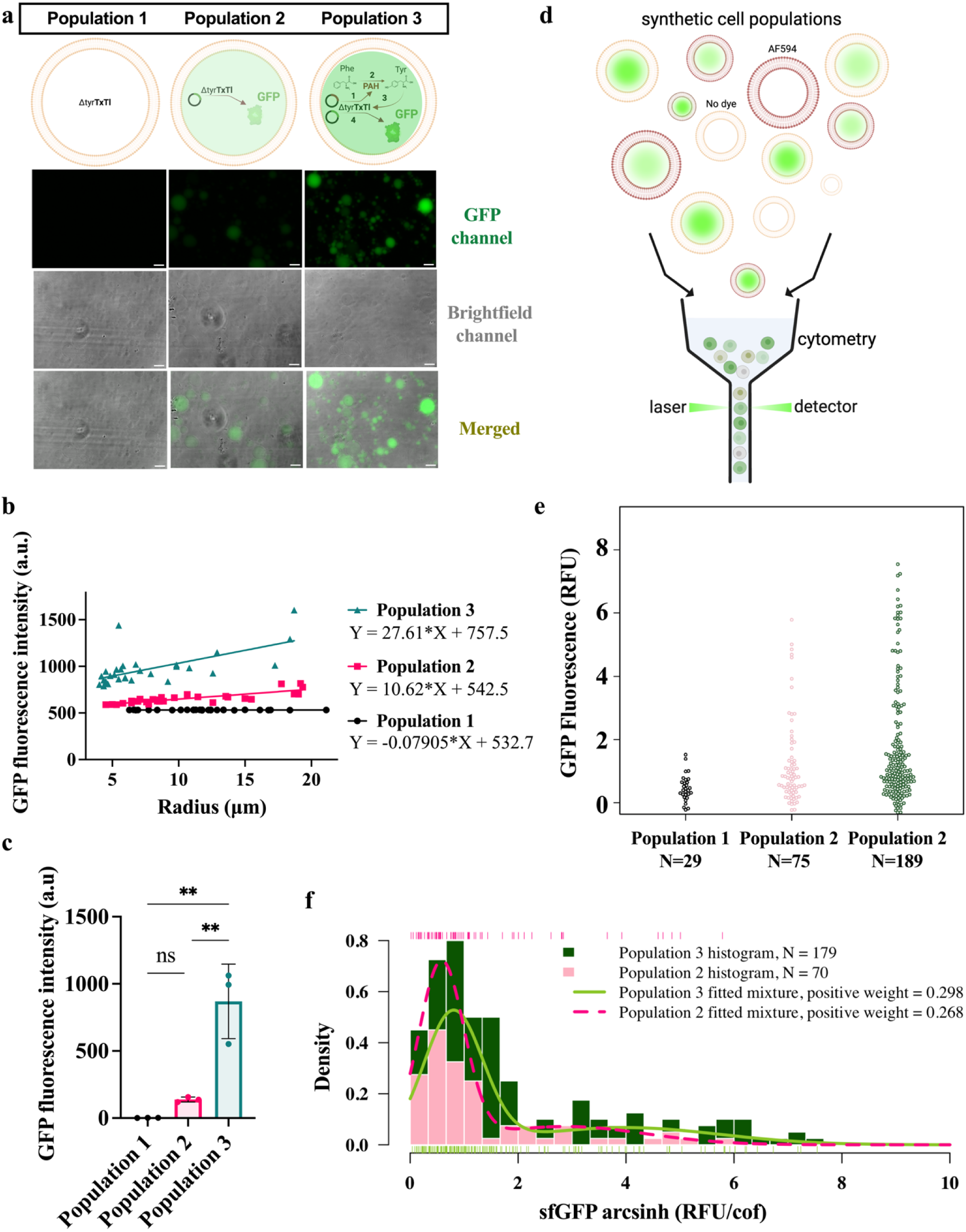
Metabolism-dependent synthetic cells were programmed using PAH-TxTl coupling. **a.** Schematic representation of three liposome populations and representative liposome images under GFP filter, brightfield, and merged. Population 1 represents no template (Δtyr-TXTL), population 2 represents background (sfGFP Δtyr-TxTl), and population 3 represents metabolically active (sfGFP and PAH Δtyr-TxTl) liposomes. In the liposomes of population 3, PAH is expressed through Δtyr-TxTl and synthesizes Tyr enzymatically when provided with substrates and cofactors. The Tyr produced by PAH then promotes sfGFP expression. Scale bar: 25 μm. **b.** Scatter plot showing mean intensities versus liposome radius for each population. Population 3 shows a higher fluorescence signal and slope, indicating that liposome size positively affects the GFP signal in this population. Analysis was done on 30 liposomes per population from raw images. **c.** ROI analysis of three liposome populations using raw images. Three images from each population were analyzed. The signal from the 10 liposomes in each image was averaged, and the background signal from the three regions of the same image was subtracted. This was repeated three times for each population. Fluorescence intensities were obtained from the mean intensities. Statistical significance in ROI analysis was assessed using a one-way ANOVA. Error bars indicate standard deviation (SD). Data is expressed as the mean ± SD. **d.** Schematic representation of flow cytometry showing that three populations, with and without the membrane dye AF594, were analyzed by flow cytometry. **e.** Distribution of sfGFP fluorescence intensities across liposome populations measured by flow cytometry. The population 2 exhibited significantly greater fluorescence than population 1 (one-sided Mann–Whitney test, *p* = 1.703 × 10□³). Population 2 showed significantly greater fluorescence than population 1 (*p* = 9.903 × 10□□) and modestly greater fluorescence than population 2 (*p* = 5.675 × 10□³). **f.** Mixture-model analysis of liposome populations. Two-component mixture models identified low-fluorescence and high-fluorescence subpopulations within populations 2 and 3. The high-fluorescence subpopulations accounted for 29.8% and 26.8% of the total flow events in populations 3 and 2, respectively. Compared to population 1, these subpopulations in population 3 and population 2 samples showed significantly greater fluorescence (one-sided Mann–Whitney tests, *p* = 8.283 × 10□¹□ and *p* = 1.014 × 10□□, respectively). The subpopulation of population 3 also exhibited significantly greater fluorescence than that of population 2 (*p* = 3.525 × 10□³).

To characterize the system more thoroughly, the same three liposome populations, with and without the membrane dye Alexa Fluor 594 (AF594), were analyzed by flow cytometry (Fig. 5d). Analysis of the entire liposome populations showed that population 3 exhibited higher maximum GFP fluorescence than both populations 1 and 2 (Supplementary Fig. 5a). Similarly, when analysis was restricted to the putative giant unilamellar vesicles in the dyed samples, population 3 again displayed the highest sfGFP fluorescence signal (Fig. 5e). To further characterize the populations, we applied two-component mixture models to populations 2 and 3 to distinguish metabolically active liposomes from lower-fluorescence events (Fig. 5f). QQ-plots supporting the model fit are shown in Supplementary Figure 6. In both populations, the models identified a low-fluorescence population that closely resembled the single population observed in population 1. These low-intensity events likely represent debris, damaged vesicles, or liposomes lacking the proper components required for cell-free transcription–translation activity. In addition to this low-fluorescence population, the models identified a second, right-skewed high-fluorescence population consistent with metabolically active liposomes. These active populations accounted for 26.8% of total flow events in population 2 and 29.8% in population 3 (Fig. 5f). It was also found that the high-fluorescence populations from both population 2 and population 3 were significantly brighter than population 1, with population 3 producing the strongest fluorescence signal overall (Fig. 5f). A comparison of active populations between population 3 and population 2 also showed a statistically significant increase in fluorescence in population 3.

These results show that synthetic cells with metabolism-dependent activity can be successfully engineered and gain a selective advantage under resource-limited conditions. Under this selective pressure, these metabolically active synthetic cells exhibit enhanced fitness relative to populations lacking metabolic function, highlighting the potential for metabolism-driven selection within synthetic cellular systems.

## Conclusion

In this work, we establish a generalizable framework for coupling genetically encoded metabolism to protein expression by engineering amino acid scarcity as a regulatory handle in cell-free systems. By selectively depleting tyrosine via dialysis and restoring it through phenylalanine hydroxylase (PAH) activity, we demonstrate that translation can be made contingent on metabolic function. This direct control of gene expression through enzyme abundance and substrate availability operates robustly across lysate-based and PURE systems, despite their distinct resource landscapes.

We extended this framework to liposome-based synthetic cells and demonstrated that even simple metabolic modules can generate distinct phenotypic populations. These populations showed measurable fitness advantages under nutrient limitation, despite intrinsic heterogeneity arising from compartmentalization.

Together, these results highlight amino acid metabolism as a powerful and tunable interface between biochemical activity and genetic output. More broadly, programming resource-aware synthetic cells will advance the development of increasingly autonomous and self-regulating biological systems.

## Acknowledgments

This work was supported by Alfred P. Sloan Foundation grant G-2024-22710 and ARPA-H award B674286 to K.P.A, NSF award 2338121 to A.E.E, Fulbright student core program/PhD, Grant number TR-SCP-2023-32 For B.V.U, and NSF award ITE-2452482 to V.N.

## Materials and Methods

### General conditions and replicates

All experiments shown in this paper were done in triplicate. All replicates for PCR, RT-qPCR, fluorometric analysis, UV-Vis analysis, cytometry, SDS-PAGE and western blot, agarose gel analysis, and sequencing were prepared from independently prepared samples. For liposome experiments, each replicate was prepared from a separate thin lipid film. Unless otherwise indicated, bulk TxTl reactions and TxTl reactions in synthetic cells were incubated at 30°C without repeated shaking. Unless otherwise indicated, PURE reactions were incubated at 37°C without repeated shaking. All experiments were performed under an RNAse-free regime, with RNAse-free equipment. All graphs were prepared and analyzed with GraphPad Prism.

### Plasmids and cloning

Two different backbones were used for in vitro expression analysis. These are p2008 with T7 max promoter and p2955, taken from Jewett lab (pJL1 plasmid) with T7 promoter. These plasmids are optimized for bacterial in vitro transcription; both contain the UTR1 region, and p2008 contains the T500 terminator, whereas p2955 contains T7 terminator sequences^22^. For cellular PAH expression, the pET32a backbone with T7 promoter and T7 terminator was used^22^. Primers were designed using the NEB Builder tool. Gibson assembly kit (NEB, E5510S) was used for assemblies, and NEB® 10-beta Electrocompetent E. coli (C3020K) was used for transformations. Colonies were seeded on agar plates and grown in lysogeny broth with a selective antibiotic marker. Plasmids were purified using GenCatch™ Plasmid DNA Mini-Prep Kit (Epoch, 2160250) and sequenced.

### AKABY cell extract preparation

The 3X AKABY cell extract protocol^23^ was adapted from Noireaux^24^ and Jewett^25^. First, a primary culture was initiated by inoculating 50 mL of LB medium supplemented with kanamycin with a loopful of cells from a glycerol stock and incubating at 30 °C for 20 hours. The culture was then held at room temperature on the bench until 7:00 PM, after which 2.5 mL of the primary culture was used to inoculate 1 L of 2×YTPG medium (2×YT broth, 31 g·L□¹; potassium phosphate monobasic, 22 mM; potassium phosphate dibasic, 40 mM; glucose, 2% w/v). The secondary culture was grown overnight until it reached an OD□□□of 0.5-0.6. Cultures were harvested by centrifugation at 3,300 rpm for 30 minutes at 4 °C using pre-cooled 750 mL bottles, and the supernatant was discarded. Cell pellets were washed twice with 200 mL cold Wash Buffer A (Tris-acetate, 10 mM, pH 8.2; magnesium acetate, 14 mM; potassium acetate, 60 mM) supplemented with freshly added DTT (2 mM final) and centrifuged at 3,500 rpm for 30 minutes at 4 °C before each wash. After the final wash, pellets were resuspended in 40 mL Wash Buffer A, transferred to pre-weighed 50 mL conical tubes, and centrifuged again at 3,500 rpm for 30 minutes at 4 °C. The wash buffer was removed, and the pellet mass was measured. Pellets were flash-frozen in liquid nitrogen and stored at -80 °C until further processing. For lysis, pellets were thawed on ice in the cold room and resuspended in Wash Buffer A containing DTT at 1.1 mL per gram of cell mass for 30 minutes, followed by gentle homogenization with a clean, cut pipette tip. Cells were lysed by sonication on ice in a 15 mL tube using 10s on/15 s off pulses at 50% amplitude until the calculated energy input was reached (2.7 × sample volume in mL / 4.5 kJ). Lysates were clarified by ultracentrifugation at 15,000 × g for 30 minutes at 4 °C, and the supernatant was collected and subjected to a runoff reaction by incubating 500 µL aliquots at 37 °C for 1 hour with tube lids slightly open. Following the runoff, samples were centrifuged again at 15,000 × g for 30 minutes at 4 °C, and the clarified supernatant was carefully transferred to new 1.7 mL tubes without disturbing the pellet. For dialysis, 2 L of Wash Buffer A supplemented with 2 mM DTT was used. The extract was loaded into a 10 kDa dialysis cassette (Thermo Fisher, 66380) using a 6 mL syringe fitted with a 1.2 mm needle, equilibrated in buffer in the cold room with gentle stirring, and dialyzed with buffer exchanges after 2 hours and again after an additional 2 hours, followed by overnight dialysis. After dialysis, the extract was recovered from the cassette. Ribosome and protein concentrations of dialyzed and non-dialyzed extracts were quantified by NanoDrop spectrophotometry at A260 and A280, respectively. If dialysis resulted in dilution, samples were reconcentrated using a Thermo Scientific Pierce 3 kDa MWCO concentrator to restore ribosome and protein levels to those of the undialyzed extract.

### Cell extract-based cell-free transcription (TX) and translation (TL)

A 10× energy mix was prepared using the previously defined protocol^20^ with following components and final concentrations of 15 mM adenosine triphosphate (ATP) and guanosine triphosphate (GTP); 9 mM cytidine triphosphate (CTP) and uridine triphosphate (UTP); 0.68 mM folinic acid; 2 mg·mL□¹ *Escherichia coli* transfer RNA (tRNA) mixture; 3.3 mM nicotinamide adenine dinucleotide (NAD); 2.6 mM coenzyme A (CoA); 15 mM spermidine; 40 mM sodium oxalate; 7.5 mM cyclic adenosine monophosphate (cAMP); 300 mM 3-phosphoglyceric acid (3-PGA) as the energy source; and 500 mM HEPES buffer adjusted to pH 8. Additionally, a 20 mM amino acid mix was prepared using the previously defined protocol^27^ by combining the amino acid powders and dissolving them in 400 mM KOH. TxTl reaction mixture contained the following components and final concentrations: 12 mM magnesium glutamate; 140 mM potassium glutamate; 1 mM dithiothreitol (DTT); 1X energy mix; 2 mM amino acids; DNA template at various concentrations; murine RNase inhibitor (40 U·µL□¹ s (50X)) at 1X; 1.5 µM T7 RNA polymerase; and cell extract (3X) at a final concentration of 1X. Whenever PAH was involved in the TxTl reaction, whether purified PAH or PAH plasmid, the water volume was replaced with PAH reaction components at a final concentration of 1X, and DTT was excluded from the TxTl mix. TxTl reactions containing the GFP template, with or without purified PAH, were carried out at 37°C for 8 hours, with real-time monitoring of the fluorescent signal using a qPCR (Bio-Rad, C1000 Touch) with a SYBR filter to reduce the detection limit and obtain a stronger readout. 96-well Hard-Shell PCR plates (Bio-Rad, HSP9601) were used for qPCR readings. Readings were taken every 5 minutes for 8 hours. TxTl reactions containing the sfGFP template, with or without PAH plasmid, were carried out at 30°C for 16 hours with real-time monitoring of the fluorescent signal using a Gemini SpectraMax microplate fluorescent reader (SpectraMax) at 485 nm excitation and 510 nm emission, and PMT sensitivity was set to low to prevent signal saturation and reads were taken with a column priority read mode for 16 hours with 20 minutes intervals. 384 well black plates(Corning, 21807021) were used for fluorescent reader-based readings.

### Protein synthesis using recombinant elements (PURE)

PURExpress (NEB, #E6840) expression was performed based on the instruction manual. Supplements include a final concentration of 0.3 mM amino acid mix and 20 U RNase inhibitor. Plasmid concentrations varied across experiments. When purified PAH or a PAH plasmid was used, the water volume was replaced with PAH reaction components, resulting in a final concentration of 1X. Reactions were performed at 37°C for at least 2 hours and 30 minutes, with real-time monitoring of the sfGFP signal using a qPCR (Bio-Rad C1000 Touch) with a SYBR filter to reduce the detection limit and obtain a stronger readout. 96-well Hard-Shell PCR plates (Bio-Rad, HSP9601) were used for qPCR readings. Readings were taken every 5 minutes for 8 hours.

### PAH expression and purification

The PAH sequence was retrieved from the UniProt database and optimized for *E. coli*^28^. A C-terminal 6XHis tag was included for purification purposes. The PAH sequence was cloned into a pET32a vector via Gibson Assembly and transformed into 10-beta electrocompetent *E. coli* cells (NEB) using electroporation. The construct was sequenced for quality control. Subsequently, it was transformed into BL21(DE3) competent cells (NEB, C2527) through heat shock. The expression and purification were performed according to the protocol described by Deich et al.^29^.

### PAH enzymatic reaction and determination of Michaelis-Menten parameters

The PAH reaction mixture contained the following components and final optimized concentrations: 0.1 M sodium HEPES buffer (pH 7.6); 4 mM dithiothreitol (DTT); 0.5 mM using the synthetic form of tetrahydrobiopterin, which is 6,7-dimethyl-5,6,7,8-tetrahydropterin (DMPH□); 2 mM L-phenylalanine; 6 µM ferrous sulfate; and 1.0 µM cPAH (test condition) or deionized water (negative control). These components were incubated at 37 °C for 3 hours, and the activity was measured using UV-vis (Agilent Cary 60 WinUV) at absorbances of 274.25 nm and 257.5 nm. Changes in amino acid concentrations were calculated using the extinction coefficients of Tyr and Phe at these wavelengths, according to the Beer-Lambert Law ^30,31^. Michaelis-Menten parameters were determined by performing the same optimized enzymatic assay at different Phe concentrations (5 mM, 0.5 mM, 0.25 mM, 0.05 mM) with and without PAH enzyme in a multicell cuvette holder. Fluorescence was measured in kinetic mode using a spectrofluorometer (Cary Eclipse). Data were collected in fluorescence mode with excitation scanning. The excitation wavelength was set to 275 nm, while emission was monitored at 303 nm. Both excitation and emission slit widths were set to 5 nm. Dwells times were set to 1 second, and readings were cycled for 4 cuvettes every 12 seconds. Measurements were taken for 5 minutes. The stage was set to 37 °C. Excitation filtering was set to automatic, and the emission filter was open. The photomultiplier tube (PMT) voltage was set to 750 V. This measurement was repeated for 3 replicates for each concentration. Tyrosine standards at concentrations of 0.5 mM, 0.25 mM, 0.125 mM, and 0 mM were detected using the same spectrofluorometer in scanning mode, with the same settings. A linear regression graph was prepared. Tyrosine concentrations from the PAH reaction were calculated by fitting fluorescence values coming from the PAH linear regression graph. To generate the Michaelis-Menten graph, initial velocities were measured, and nonlinear regression was performed at increasing Phenylalanine concentrations to estimate Km and Vmax values.

### Synthetic cell liposome preparation

Liposomes for microscopy analysis were prepared with 1mM POPC (1-palmitoyl-2-oleoyl-glycero-3-phosphocholine, Avanti 850457C) phospholipid in mineral oil (0.84g/mL)(Santa Cruz, 202539A). For cytometry, 1 mM total phospholipids were mixed with 0.3 mM cholesterol in the same mineral oil with 25 μM ABIL WE09 and 6% (v/v) n-decane. These phospholipids included POPC, DOPC (1,2-dioleoyl-sn-glycero-3-phosphocholine, Avanti 850375C), and 30mol% POPE (1-palmitoyl-2-oleoyl-sn-glycero-3-phosphoethanolamine, Avanti 850757C). For cytometry analysis, liposomes with fluorescently labeled lipid bilayers were accomplished by amending the above lipid composition with 1mol% (10uM) AF594 PE, (1,2-dioleoyl-sn-glycero-3-phosphoethanolamine-N-(TopFluor™ AF594), Avanti 810387C). To prepare a lipid-in-oil (LIO) for cytometry, a volume of mineral oil with surfactant and n-decane is measured into a graduated cylinder by mass. Using this volume of the mineral oil as the final solution volume to determine the amount of lipids to add, chloroform stock solutions of the desired lipids are added to the graduated cylinder. To evaporate the chloroform, the cylinder is placed in a fume hood, covered in foil to protect fluorescently labeled lipids, and a long needle is used to sparge the mineral oil solution with a gentle stream of nitrogen. LIO solution preparation was begun in the evening hours, and after sparging to remove the chloroform overnight, the LIO was aliquoted into ampoules. Ampoules are flushed with a gentle lamellar stream of nitrogen at 40 psi and are promptly sealed with a horizontally held Bunsen burner. Ampoules of LIO solution are stored at -20 °C, and before use, they are removed to equilibrate at room temperature to avoid opening them when condensation is present. To prepare the LIO solution for microscopy, POPC in chloroform was added to mineral oil in a glass vial, and the chloroform was evaporated at 75 °C for an hour with the vial cap open. At the end of the incubation, the mixture was allowed to cool at room temperature for 5 - 10 minutes.

Liposomes were prepared according to the emulsion transfer method described earlier, with some modifications^32^. A feeding solution (200 µL per sample as two replicates) was prepared in 1.7 mL microcentrifuge tubes. The feeding solution includes everything in the TxTl mix, except T7 polymerase, plasmids, and cell extract, which are replaced with 50% glycerol, ddH2O, and wash buffer A, respectively. TxTl reactions (10 µL per sample as two replicates) were prepared and overlaid with 200 µL of the mineral oil-lipid mixture for each replicate. The samples were emulsified using the washboard method. The emulsion was carefully transferred onto the surface of the feeding solution. Samples were centrifuged at 5,000 rcf for 10 min to pellet the liposomes. Following centrifugation, the oil phase was removed from the top of the tube. The liposome pellets from two replicates were collected from the remaining aqueous phase using a 10 µL pipette and combined with 25 µL of fresh feeding solution. Approximately 55 μL liposome solution was prepared for each sample using this method. All the liposome preparation steps were performed at room temperature. Real-time fluorescence data from TxTl reaction expressing sfGFP were collected from undiluted liposome populations for 16 hours at 30°C in qPCR, using a filter covering the excitation wavelength of 485 nM and the emission wavelength of 510 nM. Undiluted samples were used for fluorescence intensity analysis in microscopy, and ½-diluted samples were used to obtain representative images. For cytometry analysis, 20 µl of liposomes were mixed with 1ml of feeding solution.

### Fluorescent microscopy analysis

Microscopy images were captured with an Olympus IX81 inverted microscope using a 20X objective controlled by Metamorph software. Liposomes were seeded in 384-well plates (242764, Thermo Fisher). Multiple wells were imaged simultaneously in time-lapse mode. The temperature was maintained at 30 °C. Bright-field, green, and red fluorescence images were recorded using phase contrast, GFP, and Texas Red filter sets, respectively. Image processing and analysis were carried out using Fiji (ImageJ). For analysis, raw images were used. To prepare representative GFP images, the background was subtracted, all images were set to the same minimum (0) and maximum (1525) intensity levels for fair comparison, and a green LUT was applied. To prepare brightfield representative images, an unsharp mask with a radius of 1 and a weight of 0.4 was applied. To prepare merged panels, GFP and brightfield images were overlaid.

### Flow Cytometry analysis

The fluorescence intensities of Alexa Fluor 594-conjugated phospholipids and sfGFP within individual liposomes were simultaneously measured using a Cytek Aurora (Cytek Biosciences). Before measurement, the liposome dispersion was diluted with additional exogenous buffer used during liposome preparation to a final volume of 1 mL per sample. The sfGFP fluorescence was excited with a 50 mW 488 nm laser and detected through a 525(17) nm filter, while the AF594 fluorescence was excited with a 50 mW 561 nm laser and observed via a 598(20) nm filter. Since sfGFP is hypsochromic relative to AF594, the fluorescence of the membrane label is unlikely to significantly impact sfGFP fluorescence; therefore, unmixed data were used for analysis. All fluorescence intensity data were transformed using an inverse hyperbolic sine function, with scaling cofactors set at 1250 for sfGFP and 750 for AF594 to distinguish known populations in single-label and unlabeled controls. Manual gating of synthetic cells - either with or without membrane dye - was performed using OMIQ (https://www.omiq.ai/), utilizing single-label and no-label controls to identify liposome populations. For unlabeled samples, synthetic cells were identified based on forward- and UV-side-scatter areas, with side-scatter intensities transformed logarithmically to distinguish vesicles from debris. The membrane dye-labeled samples provided an alternative method for identification and gating; dyed vesicles were defined as events with the highest AF594 signal area within an acceptable UV side-scatter range to exclude debris. Finally, with both dyed and unlabeled liposomes identified, singlet events were selected for data analysis based on forward-scatter signal area and height.

### SDS-PAGE and Western blot analysis

Protein samples were mixed 1:1 (v/v) with 2× SDS-PAGE loading buffer (100 mM Tris HCl, 2.5% SDS, 20% Glycerol, 0.1% Bromophenol Blue, and fresh 4% Beta-mercaptoethanol) and denatured at 95 °C for 5 minutes SDS - PAGE was performed in a standard vertical electrophoresis system using 1X running buffer (25 mM Tris, 192 mM glycine, 3.47mM SDS) on BIO-RAD Mini PROTEAN TGX precast gels. Electrophoresis was carried out at 200 V for 30 minutes. Gels were removed and processed immediately for Western blotting or for gel fixation. For gel fixation, gels were fixed by incubation in a fixing/wash solution (10% acetic acid, 40% ethanol in water) for 30 min at room temperature with gentle rocking. After fixation, the solution was discarded, and gels were stained with Coomassie working solution (0.2% Coomassie Brilliant Blue diluted 1:1 with 20% acetic acid) for approximately 20 min at room temperature with gentle agitation. Then the staining solution was removed, and the gels were destained in the fixing solution for an hour with gentle agitation. For protein transfer, gels and membranes were assembled in a transfer cassette, and proteins were transferred at 100 V for 1 h 30 min in pre-chilled (4 °C) 1X transfer buffer (25 mM Tris, 192 mM glycine, and 10% methanol). Membranes were blocked in 5% (w/v) non-fat milk for 1 hour at room temperature on a rocking shaker, followed by incubation with primary mouse anti-His antibody (Catalog No. 652505, BioLegend) for 2 hours under gentle agitation. Membranes were washed three times with TBST (5 minutes each), then incubated with a secondary HRP goat anti-mouse antibody (Catalog No. 405306, BioLegend) and an HRP-conjugated secondary antibody for 1 hour and 30 minutes, followed by three additional TBST washes. Protein bands on gels and membranes were visualized using Pierce™ ECL Western Blotting substrate. Analysis was made based on the qualitative BLUEstain™ 2 Protein ladder, 5-245 kDa (GoldBio).

## Supplementary materials

**Supplementary Fig. 1.**
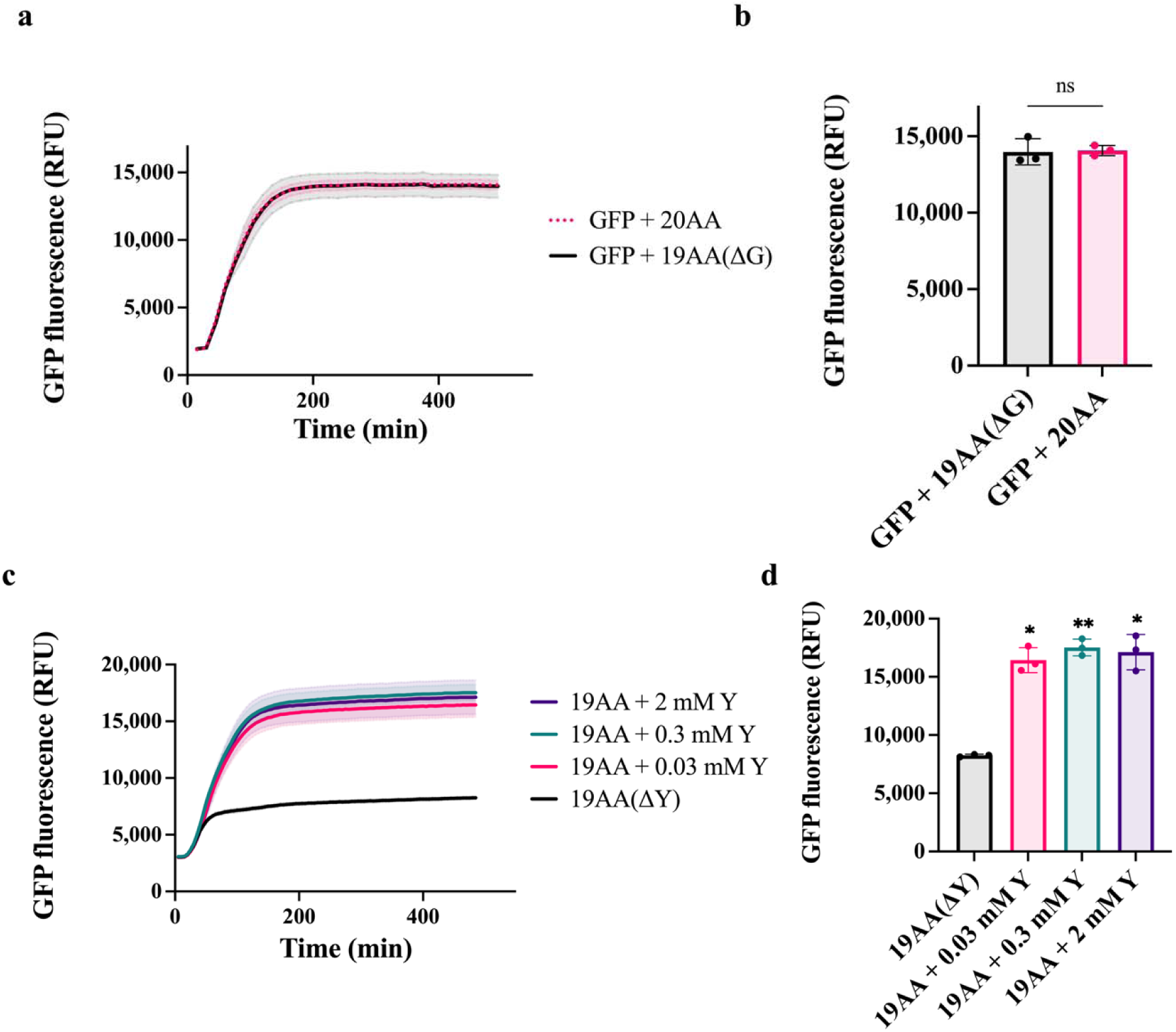
Effects of external Gly and varying external Tyr concentrations on GFP expression using. Δ**tyr-TxTl. a.** Real-time expression of GFP without external Gly. **b.** End point analysis demonstrating that external Gly supply does not improve GFP expression in Δtyr-TxTl. **c.** Real-time expression of GFP with varying external Tyr concentrations. **d.** End point analysis demonstrating that 0.03mM Tyr supply is sufficient for efficient GFP expression in Δtyr-TxTl. Statistical significance was determined using ordinary one-way ANOVA. Error bars indicate standard deviation (SD). Data is expressed as the mean ± SD.

**Supplementary Fig. 2.**
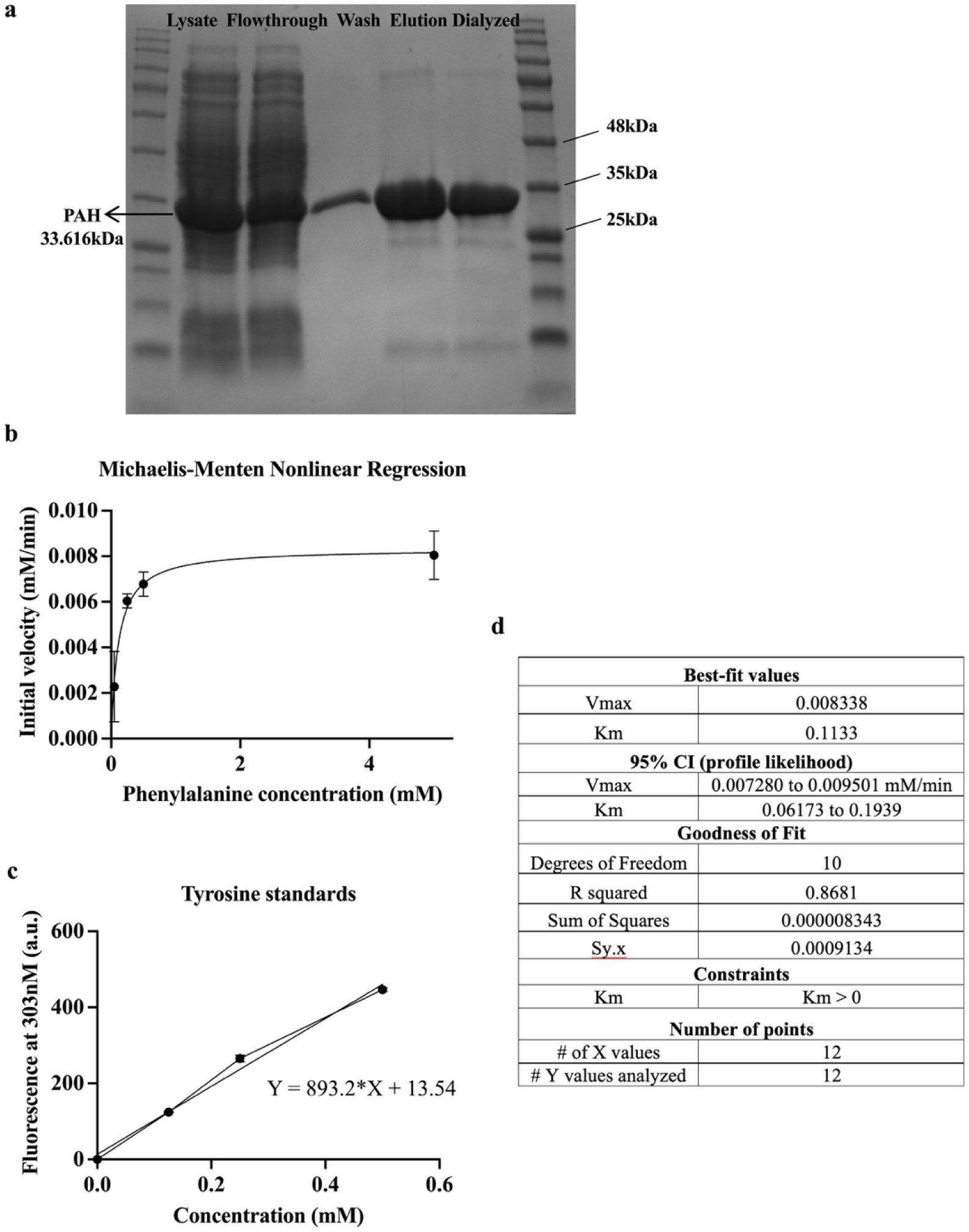
SDS PAGE analysis of Phenylalanine-4-hydroxylase(PAH) isolation from *E.coli* and kinetic parameters. **a.** SDS-PAGE analysis after PAH purification from *E. coli* demonstrates successful isolation. **b.** Michaelis-Menten graph of the PAH enzyme. **c.** Tyr standards graph was prepared to calculate Tyr produced by PAH and draw Michaelis-Menten graph. **d.** Table containing enzymatic parameters for PAH. The estimated Km (∼0.11 mM) was consistent with reported values for bacterial PAHs.

**Supplementary Fig. 3.**
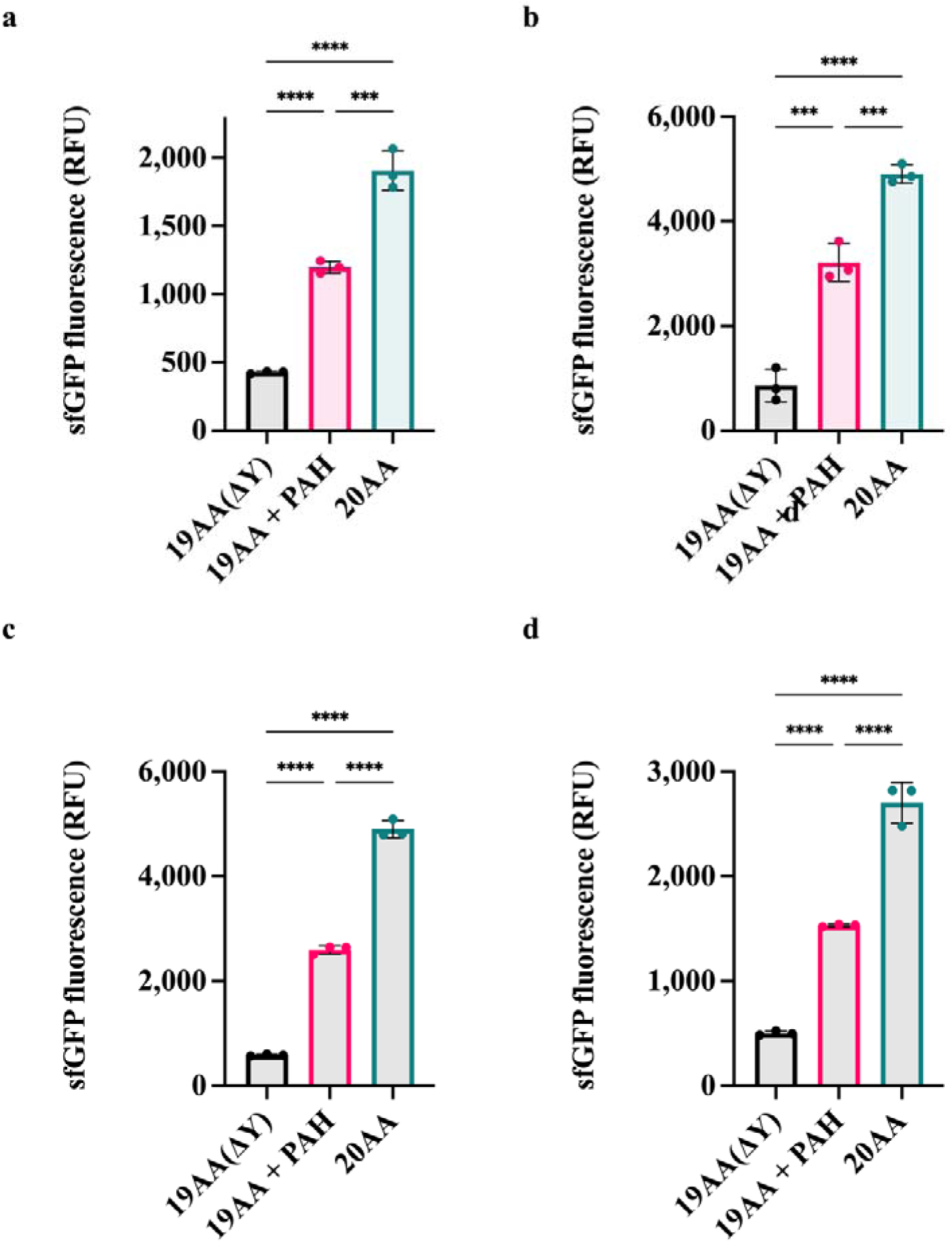
PURE expression optimization for the two plasmid system. **a.** End point fluorescence reading showing that PURE expression using 2.5 nM sfGFP and 2.5 nM PAH partially recovered sfGFP expression. **b.** End point fluorescence reading showing that PURE expression using 5 nM sfGFP and 4 nM PAH partially recovered sfGFP expression. **c.** End point fluorescence reading showing that PURE expression using 8nM PAH and 5nM sfGFP partially recovered sfGFP expression. **d.** End point fluorescence reading showing that PURE expression using 8nM PAH and 5nM sfGFP and increased substrate concentrations, Phe(4 mM), DMPH_4_ (2 mM), partially recovered sfGFP expression. Statistical significance was determined using ordinary one-way ANOVA. Error bars indicate standard deviation (SD). Data is expressed as the mean ± SD.

**Supplementary Fig. 4.**
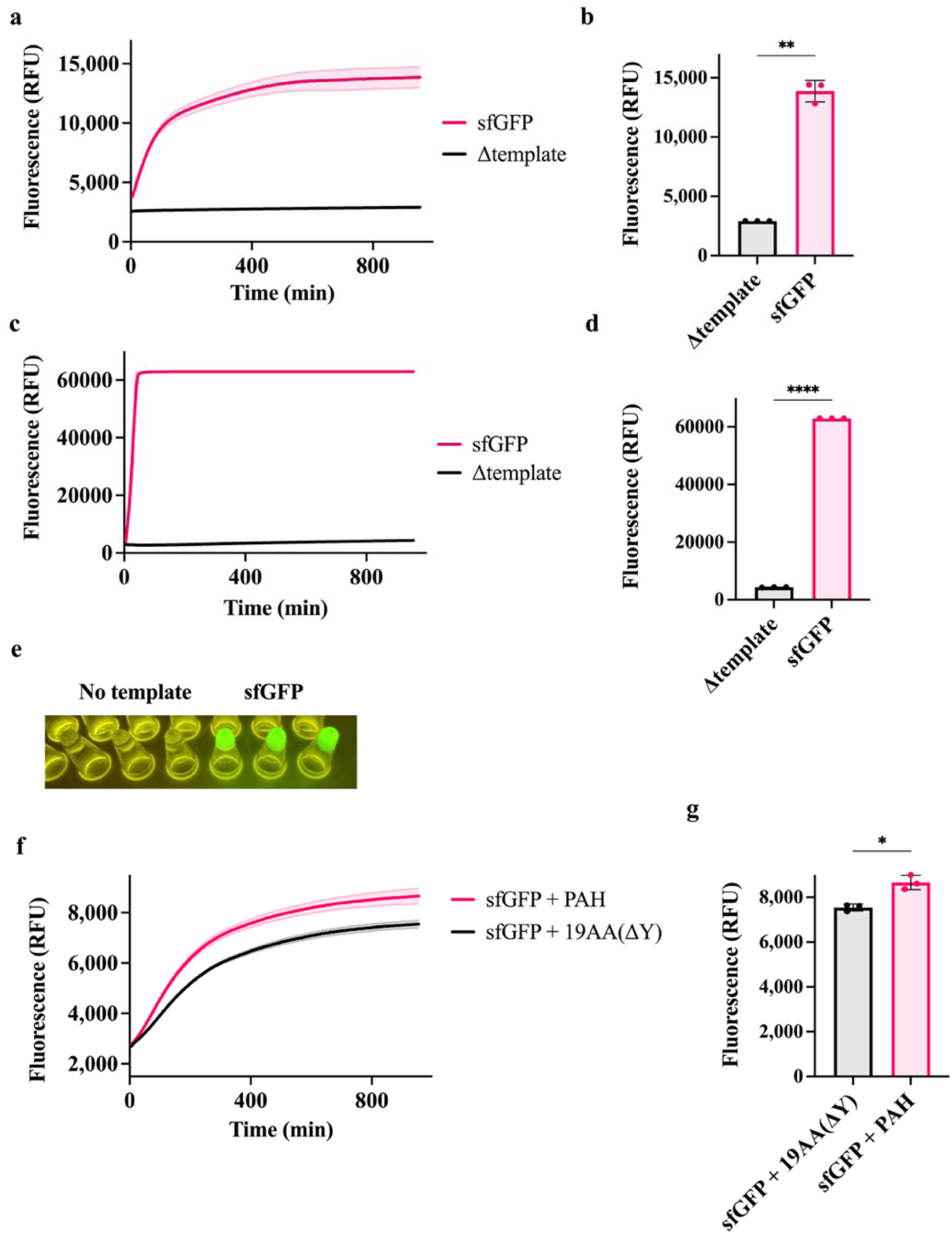
Fluorescence assessment in liposomes. **a.** TxTl containing the sfGFP template and a no-template control with 20 amino acids were encapsulated into liposomes. Real-time fluorescence readout was obtained from liposomes. **b.** End point analysis showed that the liposomes with sfGFP plasmids showed 4.2 times more fluorescence than those without template. **c.** Real-time measurement of the same TxTl reaction without encapsulation (in bulk) in real-time. **d.** End point analysis shows that TxTl yielded a 20-fold increase in sfGFP signal over the no-template control in bulk. **e.** Visual confirmation of sfGFP expression in bulk TxTl reaction. **f.** Fluorescence of liposomes encapsulating Δtyr-TxTl and PAH template was measured in a real-time assay. **g.** End point analysis showed a 1.2-fold increase in sfGFP fluorescence compared to the background control in liposomes. Statistical significance was determined using Welch’s t test. Error bars indicate standard deviation (SD). Data is expressed as the mean ± SD.

**Supplementary Fig. 5.**
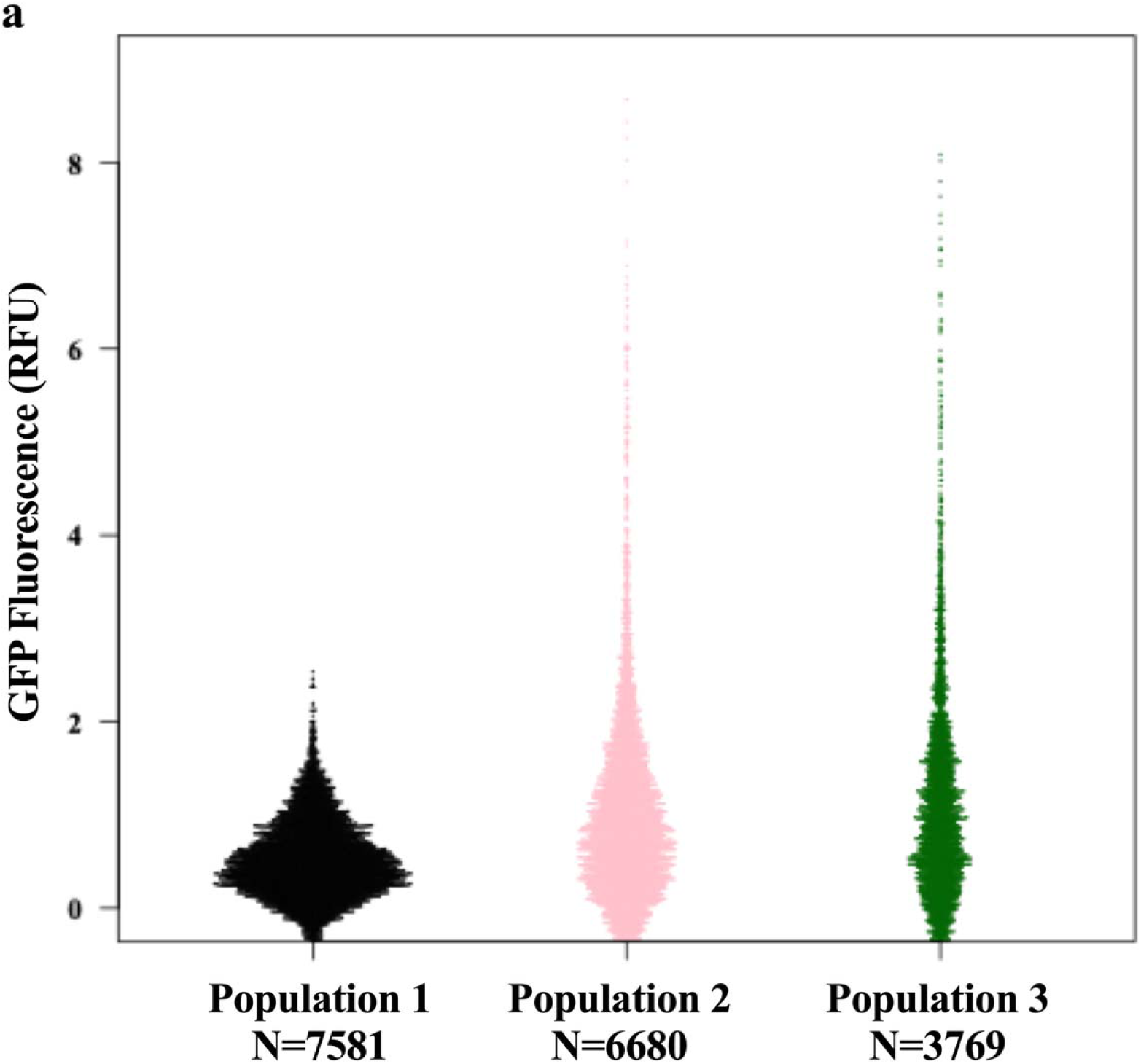
Cytometry analysis on liposome populations. **a.** Cytometry analysis of unrestricted liposome samples. The test sample showed a higher maximum sfGFP fluorescence than both the negative and baseline control populations.

**Supplementary Fig. 6.**
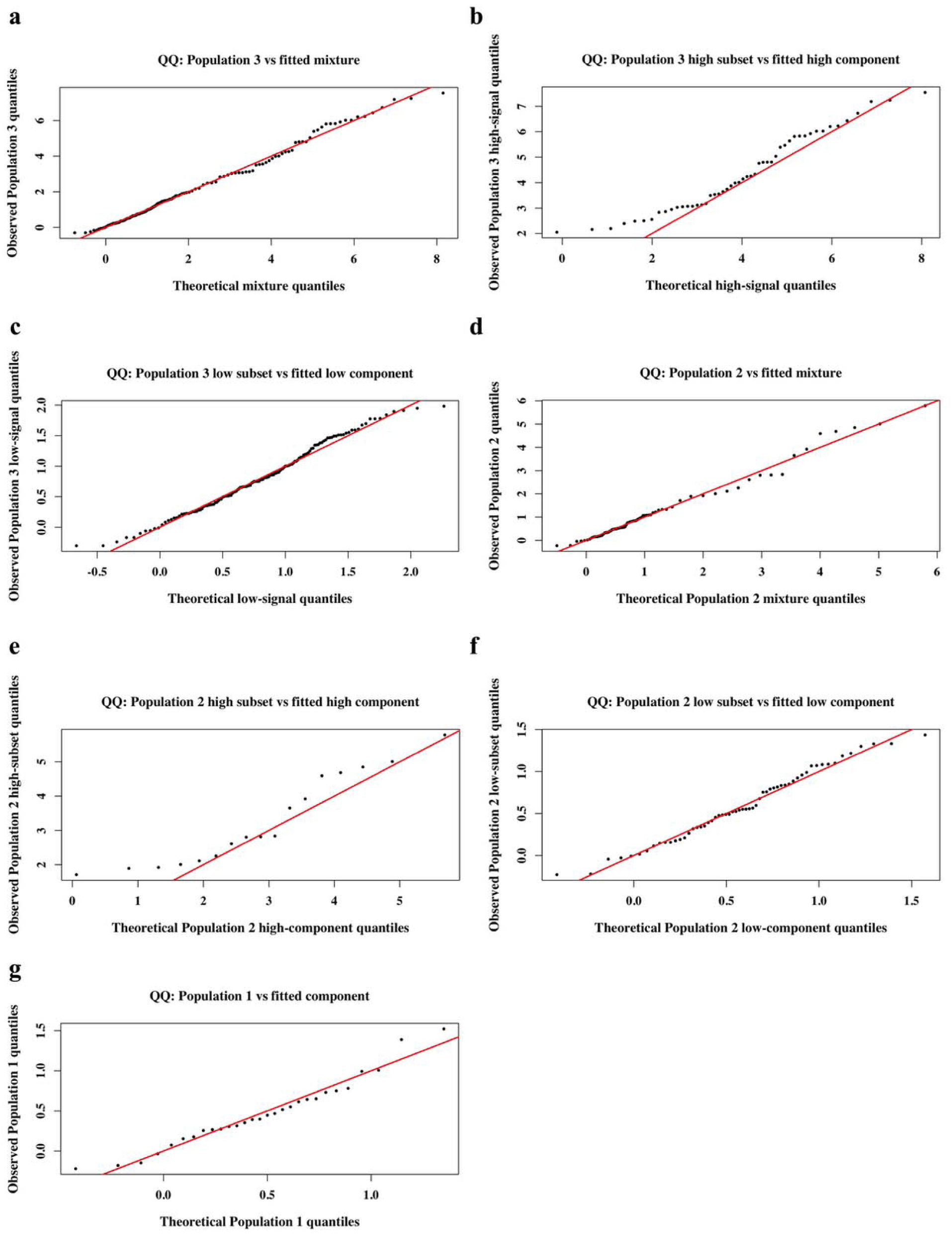
Quantile–Quantile (Q-Q) plots. QQ-plots evaluating the fit of the two-component mixture models applied to flow cytometry fluorescence distributions of liposome populations. **a.** Figure comparing population 3 vs fitted mixture. **b.** Figure comparing population 3 high subset vs fitted high component. **c.** Figure comparing population 3 low subset vs fitted low component. **d.** Figure comparing population 2 vs fitted mixture. **e.** Figure comparing population 2 high subset vs fitted high component. **f.** Figure comparing population 2 low subset vs fitted low component. **g.** Figure comparing population 1 vs fitted component.

